# Significance of the RBD mutations in the SARS-CoV-2 Omicron: from spike opening to antibody escape and cell attachment

**DOI:** 10.1101/2022.01.21.477244

**Authors:** Md Lokman Hossen, Prabin Baral, Tej Sharma, Bernard Gerstman, Prem Chapagain

## Abstract

We computationally investigated the role of the Omicron RBD mutations on its structure and interactions with ACE2. Our results suggest that, compared to the WT and Delta, the mutations in the Omicron RBD facilitate a more efficient RBD opening and ACE2 attachment. These effects, combined with antibody evasion, may contribute to its dominance over Delta. While the Omicron RBD escapes most antibodies from prior infections, epitope analysis shows that it harbors sequences with significantly improved antigenicity compared to other variants, suggesting more potent Omicron-specific neutralizing antibodies.

## Introduction

As the SARS-CoV-2 continues to spread around the world, it’s amassing mutations that occasionally lead to increased virulence and immune escape. The SARS-CoV-2 Delta variant outpaced the Alpha and Beta variants, and recently the Omicron variant, also known as B.1.1.529, is quickly taking over. Consternation about Omicron has put global health sectors on high alert due to its high transmissibility and resistance to existing therapeutic antibodies or those produced by vaccines and prior infections^1–3^. Its spread is so rapid that the cases of B.1.1.529 have been reported in >90 countries in less than a month^4^. In just three weeks since its detection in the US, it has already become the most dominant variant (>70%) followed by the Delta (^~^30%)^5^. Infections with the Delta variant are still high worldwide and though more research is needed to ascertain exactly how the presence of Omicron will affect the epidemiology of Delta, early results indicate that the immunity developed after Omicron infection is able to successfully neutralize Delta^6^. Therefore, there is a glimmer of hope that Omicron will be able to displace the more severe Delta and ultimately lessen the burden of COVID worldwide.

The large number of spike protein mutations sets the Omicron variant significantly apart from the other variants. Compared to the WT, the spike protein harbors more than 30 mutations, including 15 in the receptor binding domain (RBD) alone. The mutations in the RBD are - G339D, S371L, S373P, S375F, K417N, N440K, G446S, S477N, T478K, E484A, Q493R, G496S, Q498R, N501Y, and Y505H,^7^ compared to only 501Y in B.1.1.7, 417N, 484K, and 501Y in B.1.351, and 478K and 452R in B.1.617.2 (Fig. 1a). Specific mutations can give a variant an edge on the fitness landscape. For example, the P681R mutation in Delta is believed to have increased its transmissibility by enhancing the spike protein cleavage^8^. In the RBD, previous studies show that the mutations Q498R and N501Y increase the affinity to bind with the human receptor ACE2^9^, whereas mutations in the RBD loop region, e.g. E484K and others, are found to be associated with immune evasion^10^.

**Figure 1.**
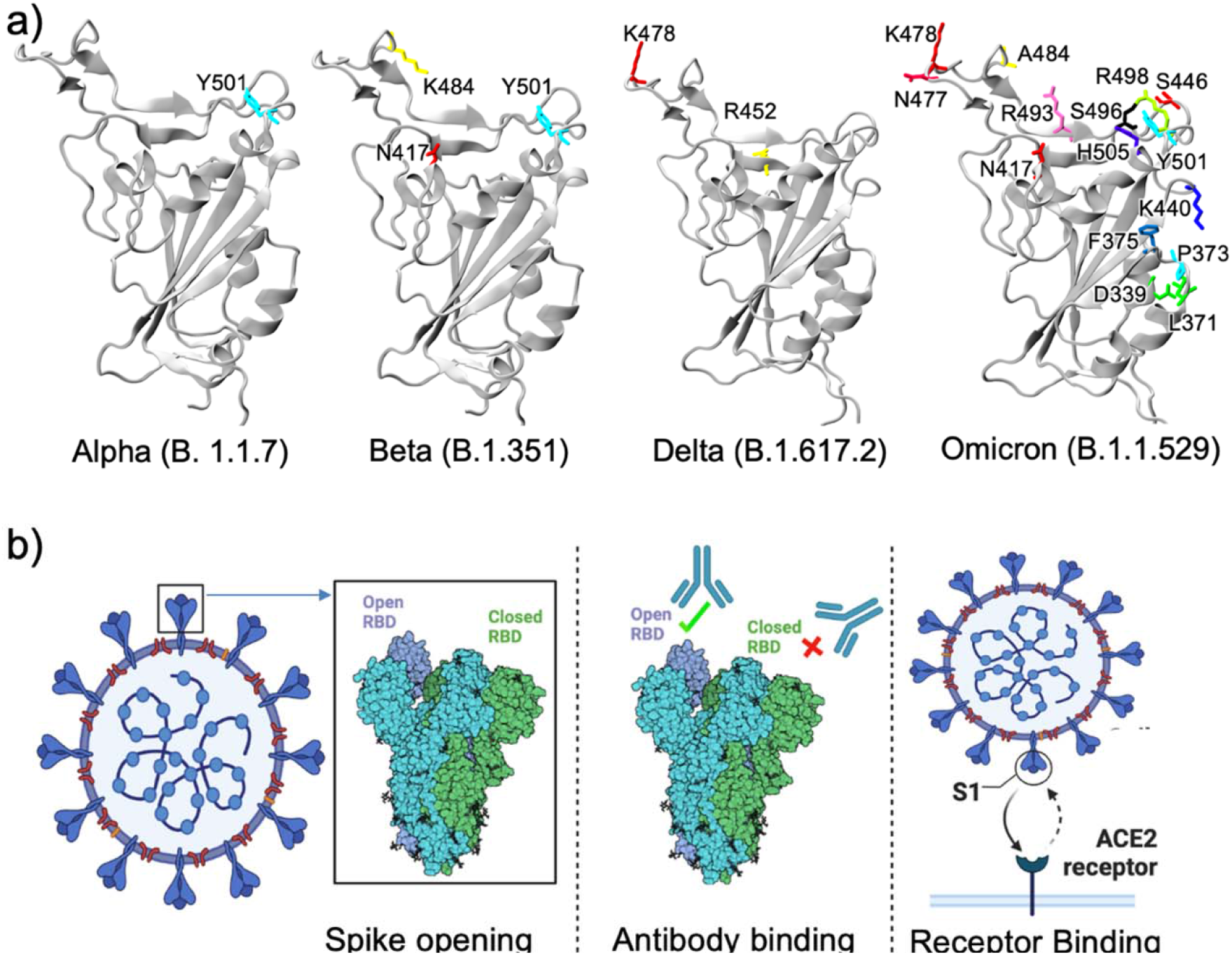
a) RBD structures of different variants. While Alpha, Beta, and Delta variants have less than three mutations in the RBD, Omicron has a remarkably large number of mutations, b) Three different ways RBD mutations may contribute to the high transmissibility of a variant (Created with BioRender.com).

Transmissibility of a respiratory viral infection such as with SARS-CoV-2 is a complex process that involves a myriad of viral, host, and environmental factors^11^. Considering only the RBD mutations, Omicron seems to have optimized its ability to infect in three different ways: 1) RBD opening 2) Antibody escape, and 3) ACE2 receptor binding. These three aspects of the RBD mutations are summarized in Fig. 1b. To investigate the consequences of the RBD mutations, we performed molecular dynamics (MD) simulations of various RBD systems for the WT, Delta, and Omicron variants. These include 1 μs MD simulations of the RBD-only system for WT and Delta, and 500 ns for Omicron, as well as 100 ns MD simulations of the RBD, together with the surrounding domains of the spike trimer, and a 100 ns simulation of the RBD-ACE2 complex. The details of the simulations, calculations, and analyses, are given in the Supporting Information, including the set-ups and systems given in Table S1.

## Results

### Effects of mutations on the RBD interactions in the closed-form trimer

The binding of the RBD with the ACE2 receptor requires the RBD to open from the closed form of the spike trimer. The three RBDs in the closed-form trimer are held together by symmetrically arranged, centrally clustered, interdomain hydrogen bonds^12^, which break during the RBD opening. In order to calculate the inter-domain hydrogen bonds formed with a RBD in a closed-form spike trimer, we prepared a simulation system for a RBD surrounded by the interacting domains of the spike trimer using the full spike trimer structure from the Amaro Lab^13^. To minimize the computational time, we prepared a truncated trimer including only the domains that directly interact with one of the RBDs (here, we chose the RBD of chain A), as shown in Fig. 2 and Fig. S1. These domains include the residues 330-530 of chain A (RBD), 16-530 and 968-1000 of chain B, and 330-530 and 968-1000 of chain C. The domains surrounding the RBD of chain A are harmonically restrained by applying harmonic forces to all C_α_ atoms that are >12 Å away from the RBD of chain A. This allows flexibility of the RBD in the trimer but maintains the integrity of the domains mimicking the full trimer. We performed 100 ns simulations for the WT and Omicron and calculated the hydrogen bonds for the last 50 ns of the trajectories. To confirm that the RBD hydrogen bonding is adequately represented by the truncated system, we calculated and compared the % hydrogen bonds for the closed-form RBD to the simulation of the full system performed by the Amaro Lab^13^. The RBD hydrogen bonding pattern has good agreement between the truncated trimer and the full trimer.

**Figure 2.**
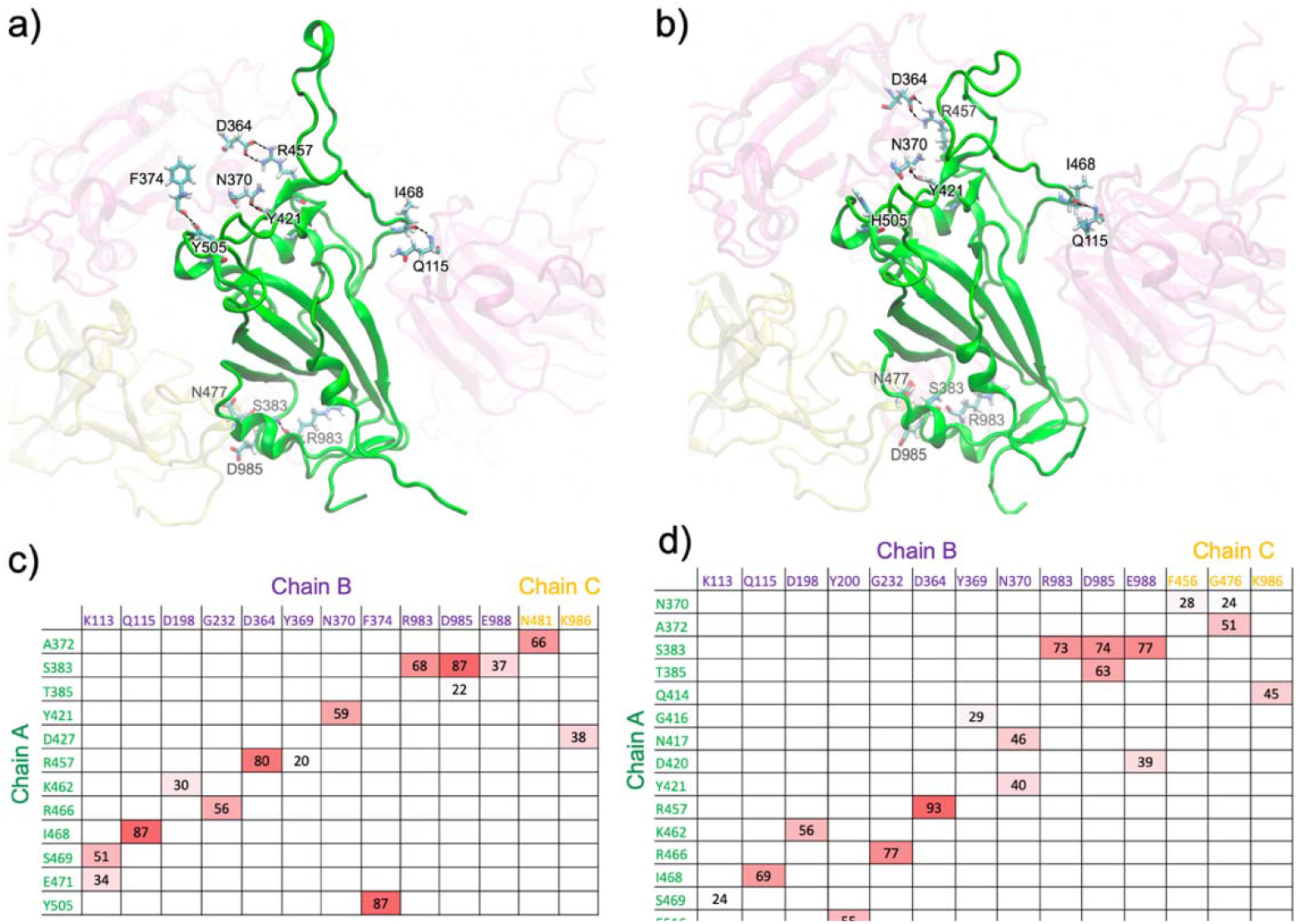
Major hydrogen bonds formed between the RBD of chain A (green) and the surrounding domains in the closed-form spike trimer for a) WT and b) Omicron. Additional interactions are shown in Fig. S1 (from different views). Hydrogen-bond pairs and % occupancies for the c) WT and d) Omicron, with the color scale from red (maximum) to white (minimum).

We show in Fig. 2 the major hydrogen bonds that an RBD forms with its surrounding domains for both the WT and Omicron. The % occupancies for the major hydrogen bonds for both the WT and Omicron are given in the matrices in Fig. 2c-d. Major hydrogen bond interactions with %occupancy >80% include R457(A)-D364(B), Y505(A)-F374(B), S383(A)-D985(B), and I468(A)-Q115(B). Residue S383 of the RBD makes three stable hydrogen bonds with the helix domain comprised of residues 968-1000 of chain B. However, only a few, weak interactions are observed between the RBD and the helix domain of chain C. The helix domains lie just below the RBD, as shown in Fig. 2 and Fig. S1. We also calculated the %hydrogen bonds (Fig. 2d) for the Omicron RBD displayed in Fig. 2b. Most of the major hydrogen bonds found in the WT are also present in Omicron, including R457(A)-D364(B), S383(A)-R983(B), S383(A)-D985(B), S383(A)-E988(B), I468(A)-Q115(B), K462(A)-D198(B), and E516(A)-Y200(B). However, one of the major WT hydrogen bonds with nearly 90% occupancy, Y505(A)-F374(B), is lost in Omicron due to the Y505H mutation. In addition, the polar to hydrophobic mutation S371L also abrogates minor hydrogen-bonding with multiple residues. When the Y505(A)-F374(B) hydrogen bond is broken, the RBD slightly repositions to make relatively weaker but new hydrogen bonds, mostly with Chain C residues. The new interactions in Omicron include N370(A)-F456(C), N370(A)-G476(C), A372(A)-G476(C), Q414(A)-K986(C), and D420(A)-E988(B). Although this loss of these hydrogen bond interactions in Omicron appears to be compensated by the increase in the %hydrogen bond or new hydrogen bonds. Therefore, it’s difficult to assess the stability of the RBD based on the total %hydrogen bonds alone. However, the opening from the closed form trimer may still be affected for the following reasons. With the Y505(A)-F374(B) hydrogen bond in the WT, the RBD of chain A is held in a position slightly away from these residues, which are in the RBD and the helix domain of chain C. These new interactions can still form in the WT if the Y505(A)-F374(B) hydrogen bond is broken and vice-versa. Therefore, our analysis shows that compared to the Omicron RBD, the WT RBD is more protected from opening from the closed-form trimer due to the possibility of either being held by Y505(A)-F374(B) or the additional interactions with chain C, whereas the Omicron RBD lacks the Y505(A)-F374(B) interaction. We note that the interactions of RBD with the neighboring domains are transient and may change as the RBD shifts. While we investigated the interactions in the closed-form state, further work is needed to determine exactly how these mutations affect the RBD opening along the opening pathway.

### Structural changes, antibody binding, and antigenic shift in the Omicron RBD

Once the RBD springs out from the closed form, it is vulnerable to antibody detection and binding due to the loss of shielding by glycans^13^. Due to mutation-induced changes in both the residue type as well as the RBD structure, antibodies elicited with prior infections or vaccines may not be able to optimally bind to the RBD. In an earlier work, we showed that the changes in the Delta RBD structure, including in the receptor-binding motif (RBM) loop segment, cause some antibodies to be ineffective^14^ at binding the RBD. With the significant number of mutations in the Omicron RBD, such effects can be extensive.

To explore the structural changes in the Omicron RBD, we performed 1.0 simulation of the RBD-only system for the WT (PDB ID 6VSB) and 0.5 Omicron (PDB ID 7T9L). We find significant differences in the RBD structure for the isolated RBD (i.e. not complexed with ACE2). Specifically, the motif consisting of residues 364 to 375, which contains the mutations S371L, S373P, and S375F, shows an extensive structural change (Fig. 3a). All these three mutations are from polar to hydrophobic and this causes the motif to realign and make non-specific interactions with F342, A435, and W436 in the hydrophobic pocket (Fig. S2 and Movie S1). As shown in Fig. 3a, the distance between the C_α_ atoms of residues 371 and 375 in the motif is relatively stable in WT, whereas it separates significantly in Omicron. Both of these residues are binding sites for antibodies (e.g. RBD-Ab complexes 7KN5 and 7M7B). This separation in the antibody-binding region can reduce or abolish the binding of the antibodies specific to these sites. Subtle structure changes in other sites in the RBD may also affect antibody binding, allowing the Omicron RBD to escape antibody detection. Figure 3b shows the antibody-binding sites identified from the RBD-Antibody complexes available in the Protein Data Bank. Almost all mutations in the Omicron RBD are located in important antibody-binding sites (residues indicated by purple boxes) and therefore can directly affect the binding of antibodies specific to the WT and other variants.

**Figure 3.**
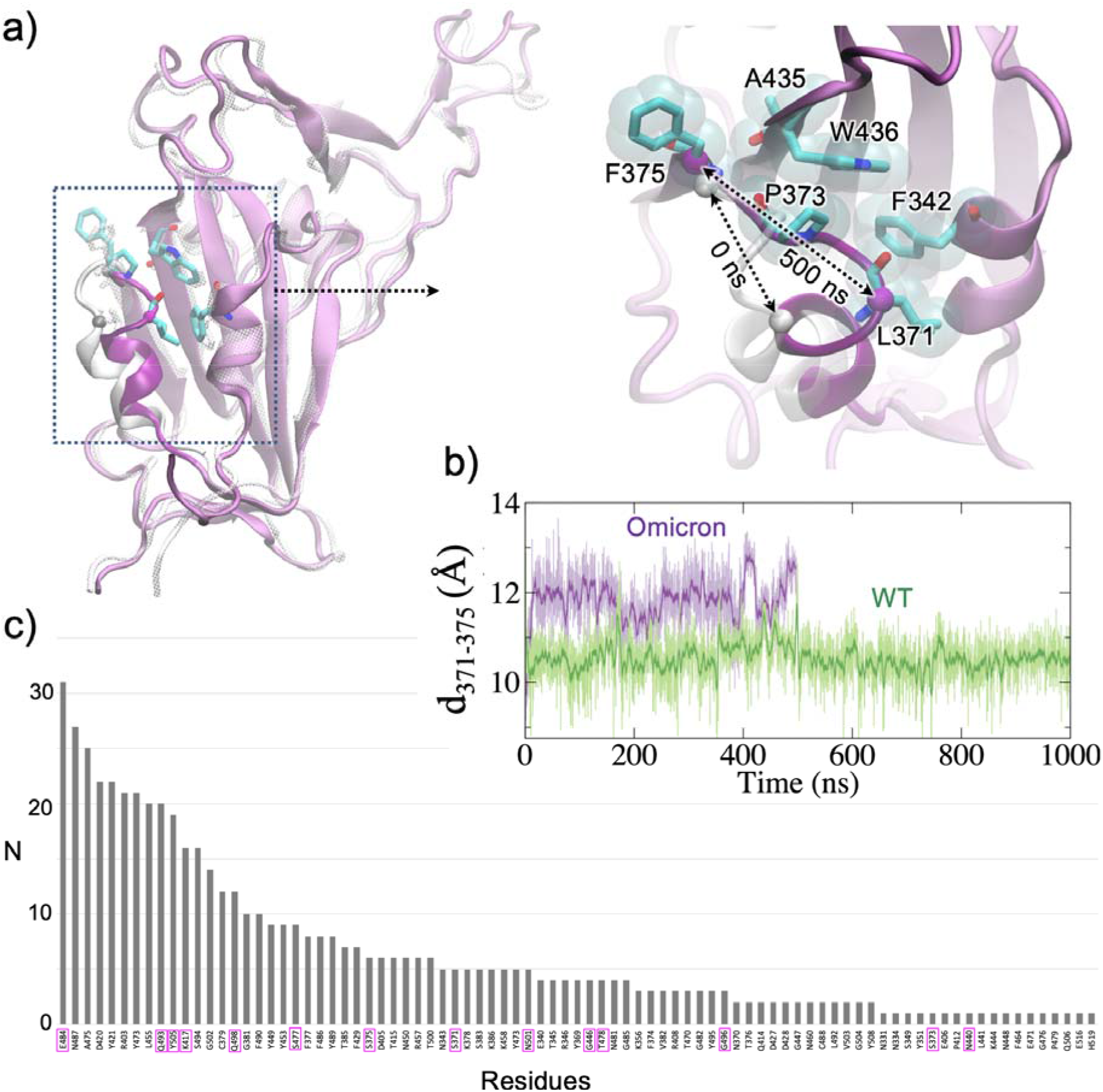
a) Structural changes in the Omicron RBD motif containing the mutations S371L, S373P, and S375F (motif highlighted in bright purple), which form a hydrophobic cluster (right). b) The C_α_-C_α_ distance between residues 371 and 375 showing the difference in WT vs Omicron. c) Number of times the RBD residues found to hydrogen bond with the antibodies in 105 RBD-Ab complexes from the Protein Data Bank. The mutated residues in Omicron are highlighted in purple boxes along the x-axis.

To explore the antigenic shifts due to the mutations, we first identified the RBD epitopes using various MHC-I and MHC-II prediction methods as well as sequence and structure-based B-Cell epitope prediction methods as described in the Supporting Information and used a consensus approach^15^ to select the epitopes for further analysis. The consensus epitopes that contain the mutations in RBD are given in the Supplementary Table S2, and all of the predicted epitopes (sequence-based and structure-based) are listed in Table S3, S4, S5 and S6. We calculated the antigenicity of the mutated sequences using VaxiJen^16^ and compared with the corresponding WT sequence to assess the antigenic characteristic introduced by the mutation. Most of the epitopes listed in Table S2 have similar antigenicity after mutations in Alpha, Beta, Delta, or Omicron. However, three epitopes in the Omicron (E2, E3, and E9 in Table S2) are found to have significantly increased antigenicity compared to the WT. These Omicron epitopes include 370-NLAPFFTFK-378 involving the mutations S371L, S373P, and S375F, 372-APFFTFKCY-380 involving the mutations S373P and S375F, and 483-VAGFNCYFPLR-493 involving the mutations E484A and Q493R. The antigenicity of E2 is 1.34 for Omicron vs 0.12 for WT. A similar increase is observed for epitope E3, which has overlap with E2. Similarly, the antigenicity of E9 is 1.23 for Omicron vs 0.56 for WT. The locations of these epitopes are shown in Fig. S3. While the mutation-induced antigenic shifts render reduced sensitivity for the WT-specific antibodies, the increased antigenicity in the Omicron epitopes suggests a more potent immune response from these epitopes.

### Effects of the mutations on ACE2-binding of the Omicron RBD

The mutations in the RBM region can directly affect the binding affinity of the RBD to bind ACE2. Kim *et al* showed differences in the force required to dissociate the RBD from ACE2 for different variants of concern (Alpha, Beta, Gamma, and Delta)^17^. With 10 mutations in the RBM region of Omicron, the effects on ACE2 attachment can be significant compared to other variants. Early results by Wu et al. suggested that Omicron RBD-ACE2 interactions is weaker than in Delta^18^. However, recent cryo-EM structure-based analysis of the RBD-ACE2 complexes of both the Omicron and Delta variants show that the Omicron RBD-ACE2 interface has better optimized interactions compared to that of Delta^19^. To investigate the effects of the Omicron mutations on the RBD attachment to the ACE2 receptor, we performed an MD simulation of the Omicron RBD-ACE2 complex (PDB ID: 7T9L)^20^ and compared the RBD-ACE2 interfacial interactions with that of WT (PDB: 6VW1)^21^ and Delta (PDB ID: 7V8B)^22^. The short MD simulations provide the dynamic nature of the interactions and allow us to calculate the probability (% occupancy) of each hydrogen bond. As shown in Fig. 4a-d, the occupancy of the inter-protein hydrogen bond in Omicron is noticeably higher than in Delta suggesting a much stronger ACE2-binding in Omicron. The major RBD-ACE2 hydrogen-bond pairs in Omicron with >70% occupancy include Y453-H34, G502-K353, N487-Y83, T500-D355, and R493-E35, and Y449-D38 with >50% occupancy. In contrast, Delta has only one interaction (K417-D30) with >70% occupancy and another (G502-K353) with 60% occupancy, suggesting a much weaker interfacial interactions in Delta compared to Omicron. This is consistent with a recent work by Genovese et al.^23^ that analyzed the interfacial interactions in the RBD-ACE2 complex using ab-initio and quantum mechanical calculations. Lupala et al.^24^ also found a significantly increased binding affinity of the Omicron RBD to ACE2, compared to the Delta RBD, suggesting an increase in infectivity. We compared the RBD-ACE2 hydrogen bonding with that in WT as reported in our earlier work^25^ (Fig. S4), and we observed additional unique hydrogen bonds (e.g. Y449-D38) in Omicron compared to WT, whereas Delta showed no unique hydrogen bonds.

**Figure 4.**
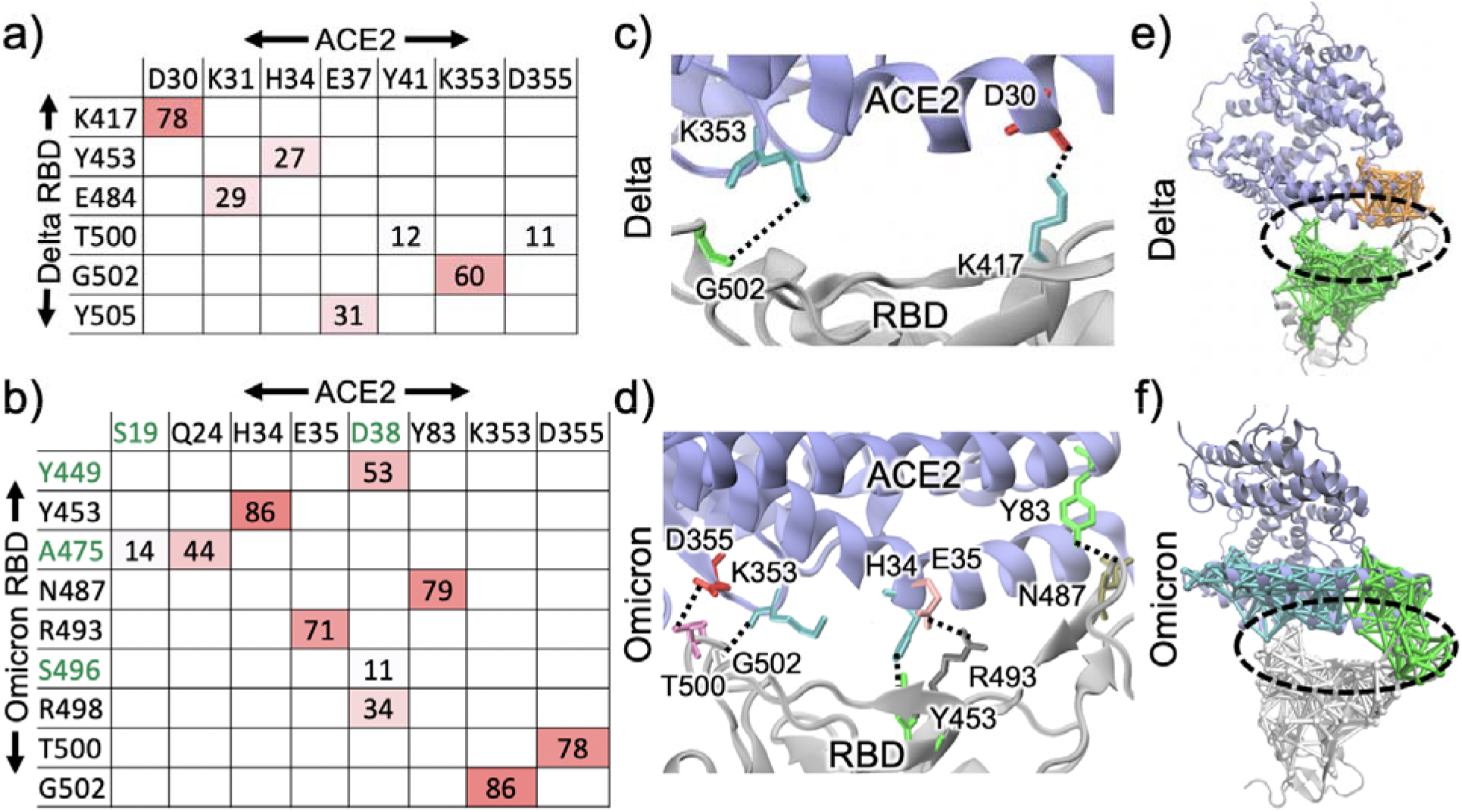
Percent occupancies of the hydrogen bonds between the RBD and ACE2 for the a) Delta and b) Omicron variants. The Unique interfacial hydrogen-bonds found in Omicron are highlighted in Green. Hydrogen bonds with >50% occupancy are shown for c) Delta and d) Omicron. The communities that span both the RBD and ACE2 are shown for the e) Delta and f) Omicron RBD-ACE2 complexes.

In addition to the increased hydrogen-bonding at the interface, the stability of the complex is also displayed by the dynamic network analysis^26^. We used the first 50 ns of the trajectory for RBD-ACE2 complexes of the Delta and Omicron variants to calculate the dynamic network communities as shown in Figure S5. We performed the community analysis by partitioning the network into subnetworks (community). A community is a collection of nodes (amino acids) that are connected with edges (connections). The communities that span across the RBD-ACE2 interface are shown in Fig. 4e,f and the corresponding residues are shown in Fig. S5b. There are a total of 23 connections that occur between the RBD and ACE2 in Omicron, compared to 7 in Delta. This suggests a better binding of the RBD to ACE2 with increased interactions that stabilize the RBD-ACE2 complex in Omicron.

## Conclusions

Overall, our analyses show that the Omicron RBD shows a higher ACE2 binding affinity compared to the Delta RBD. Unlike other variants, Omicron has RBD mutations Y505H and S371L that lie in the region interacting with the surrounding domains in the closed-form spike trimer and this can affect the RBD opening. Increased ACE2 affinity and potentially easier and more efficient RBD opening, combined with antibody evasion due to mutation-induced antigenic shifts provide the Omicron strain with a significant increase in the probability for successful cellular attachment and this may contribute to its dominance over Delta. While the Omicron RBD escapes most antibodies specific to other variants, it harbors sequences with significantly improved antigenicity compared to prior sequences. This suggests a possibility of superior neutralizing antibodies for Omicron and provides insights into vaccine design as well as a perspective on the future of SARS-CoV-2 persistence.

## Acknowledgements

This work was partially supported by the National Science Foundation under Grant No. 2037374. PB acknowledges the Dissertation Year Fellowship support from the University Graduate School at Florida International University. The authors thank the COVID-Informatics research team at Florida International University for helpful discussions.

