## Supporting Information for "Significance of the RBD mutations in the SARS-CoV-2 Omicron: from spike opening to antibody escape and cell attachment"

**Methods**

*System preparation*

For the RBD interactions with the surrounding domains in the closed-form trimer, the initial frame of the simulation trajectory of the full spike protein trimer from the COVID-19 Data Sets of Amaro lab^1^ was used. For the Omicron variant, the following mutations were introduced to the WT RBD: G339D, S371L, S373P, S375F, K417N, N440K, G446S, S477N, T478K, E484A, Q493R, G496S, Q498R, N501Y, Y505H to the RBD region (residues 330-530). For the Omicron RBD-ACE2 system, a recently deposited cryo-EM structure (PDB ID 7T9L)^2^ was obtained from the Protein Data Bank. Similarly, the cryo-EM structure (PDB ID 7V8B) was used for the Delta RBD-ACE2 simulations. The RBD-only system for Omicron was prepared with the RBD from the RBD-ACE2 complex (PDB ID 7T9L)^2^. The simulation system was set up with the same procedure as in our earlier studies^3, 4^. All the systems were prepared using the solution builder interface of CHARMM-GUI website^5, 6^.

*Molecular Dynamics (MD) Simulations*

We performed molecular dynamics (MD) simulation following the procedures used in our previous work^4^. Briefly, constant pressure MD simulations were performed with NAMD 2.14^7^ using CHARMM36 force-field^8, 9^. Following 10,000-step minimization and 2 ns equilibrations, the production runs were performed at 303.15K temperature and 1 atm pressure using 2fs timestep, and using Particle Mesh Ewald method to treat electrostatics^10, 11^ and SHAKE algorithm to constrain covalent bonds involving hydrogen atoms^12^. Visual Molecular Dynamics (VMD)^13^ 1.93 was used for the structure and trajectory visualization, as well as for the hydrogen-bond analysis. The information about the simulation systems and simulation lengths are given in Table S1. Hydrogen bond analysis was performed for the RBD-ACE2 complexes for the 100 ns of the trajectories using a cut-off of 3.5 Å and 30^o^.

*Epitope predictions*

The Spike protein sequence was retrieved from the NCBI (GenBank: QHD43416.1, accession MN908947.3), which is based upon the 2019-nCoV sequence^14^. Various independent methods were used for each of the MHC-I, MHC-II and B-cell epitopes : ProPred-I, CTLPred, and NetCTL1.2 for the prediction of MHC-I T-cell epitopes^15-17^; NetMHCII2.3 and EpiTOP3.0 for MHC-II T-cell epitopes^18, 19^; and BepiPred and BcePred^20, 21^ for the B-cell epitopes. The T-cell epitope predictions are based on quantitative matrix and Quantitative Structure-Activity Relationship models (QSAR)^18^, whereas the B-cell prediction method BepiPred uses a random forest algorithm trained on epitopes annotated from antibody-antigen structures and BcePred locates the epitopes using physiological properties such as hydrophilicity, polarity, and exposed surface. In addition to the sequence-based epitope predictions, we also used the spike trimer structure for the structure-based prediction using Ellipro and DiscoTope. Ellipro incorporates the properties such as antigenicity, solvent accessibility, and flexibility of the protein structures to predict linear and conformational epitopes. DiscoTope uses the spatial information, surface accessibility, and statistics of amino acids on the protein 3D structure to predict residue by residue conformational epitopes^22, 23^. The epitopes predicted by various methods are given in Table S3, S4, and S5. The epitopes that involve mutations in the variants of concern are summarized in Table S2 and their antigenicity was predicted using VaxiJen^24^.

Table S1. Systems simulated in this work. The WT and Delta simulations of RBD-only systems in our earlier work (Baral et. al, 2021)^25^ were extended from 600ns to 1000ns.

| System | Variant | # Of Atoms (rounded to 1000) | Box Size (Å)^3^ | Simulation Time (ns) |
| --- | --- | --- | --- | --- |
| RBD-only | WT | 78,000 | 95x95x95 | 1000 |
|  | Delta | 84,000 | 95x95x95 | 1000 |
|  | Omicron | 84,000 | 95x95x95 | 500 |
| RBD-hACE2 | Delta | 242,000 | 138x138x138 | 100 |
|  | Omicron | 254,000 | 140x140x140 | 100 |
| Closed-RBD (Truncated) | WT | 275,000 | 144x144x144 | 100 |
|  | Omicron | 275,000 | 144x144x144 | 100 |

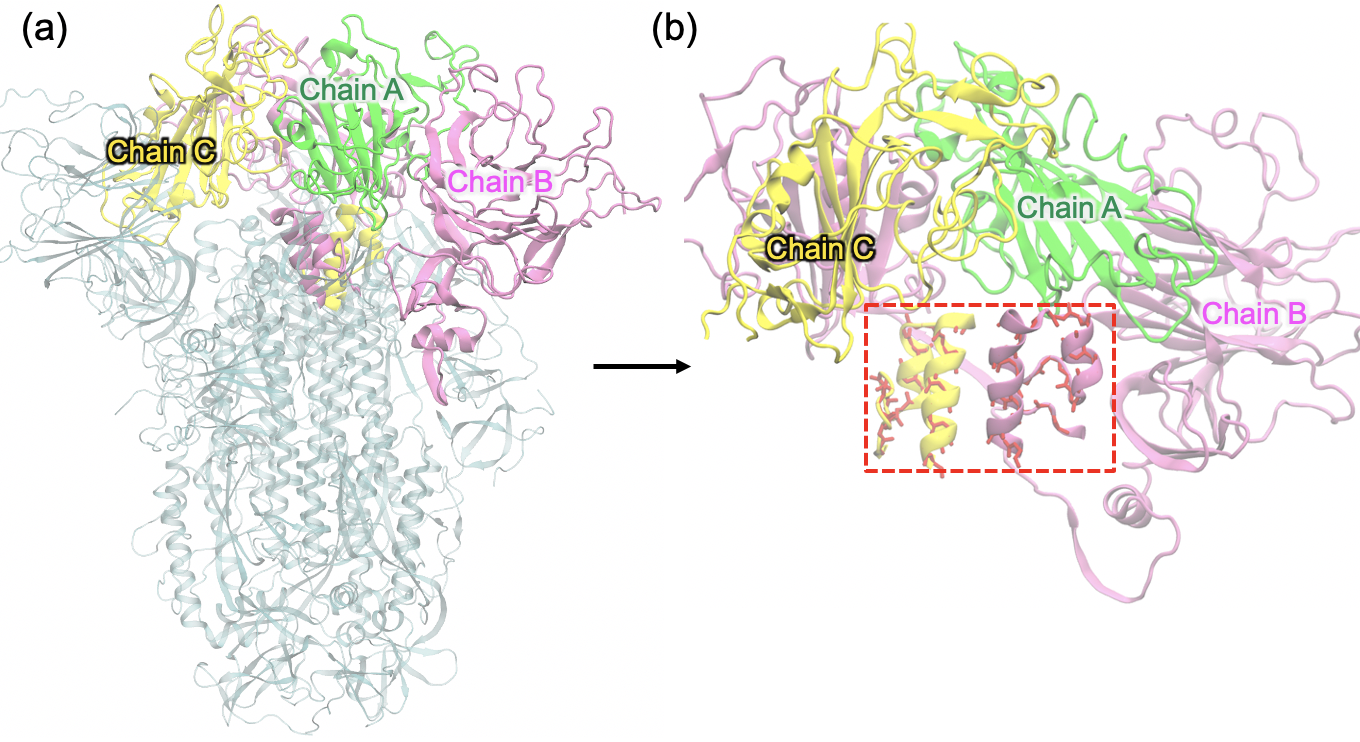

Figure S1. a) The spike protein trimer of SARS-CoV-2 in its closed form. The pdb file was obtained the first frame of the trajectory from the Amaro Lab^26^. The RBD of chain A (green) and the surrounding domains (chain B – magenta and chain C – yellow) are highlighted. b) The colored part shows the truncated system consisting of the Chain A RBD and the surrounding domains considered for MD simulations. All Cα atoms >12 Å from the RBD of chain A are harmonically restrained for MD simulations. The helical segments highlighted in red dashed box are the non-contiguous segments of chain B and C that interact with the chain A RBD. For the Omicron system, all mutations within 12 Å of RBD of chain A and the surrounding were considered.

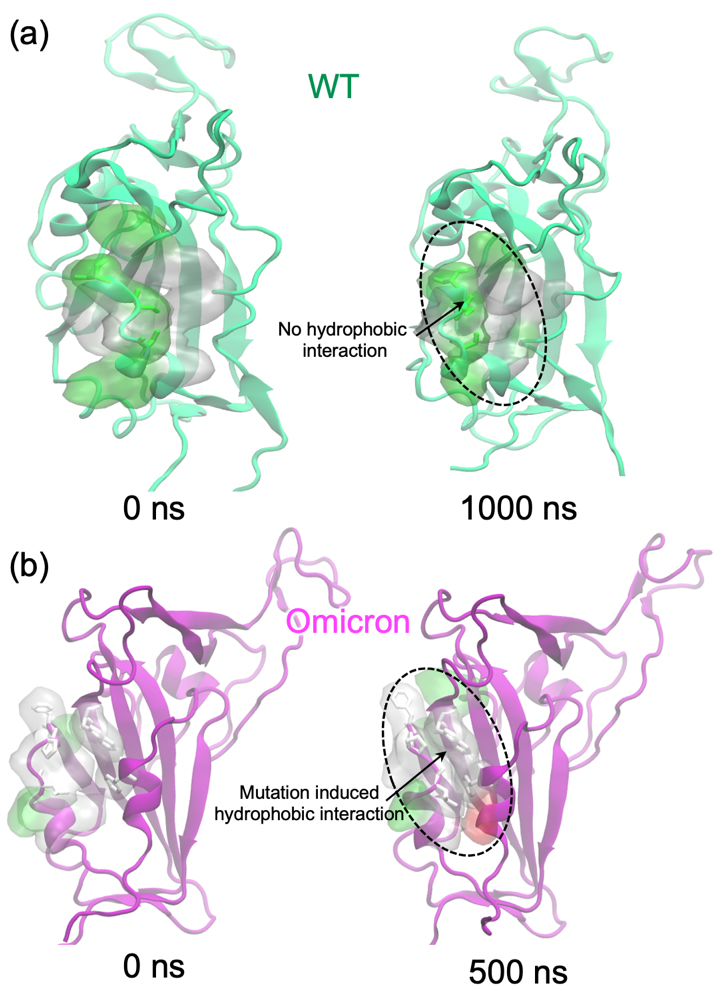

Figure S2. Comparison of the motif structure (residues 364-375) at 0 ns and 500ns for the a) WT and b) Omicron RBDs. The polar to hydrophobic mutations S371L, S373P, S375F in Omicron allow interactions with the nearby hydrophobic residues. The polar residues are highlighted in green surface and the hydrophobic residues are highlighted in white/gray surface.

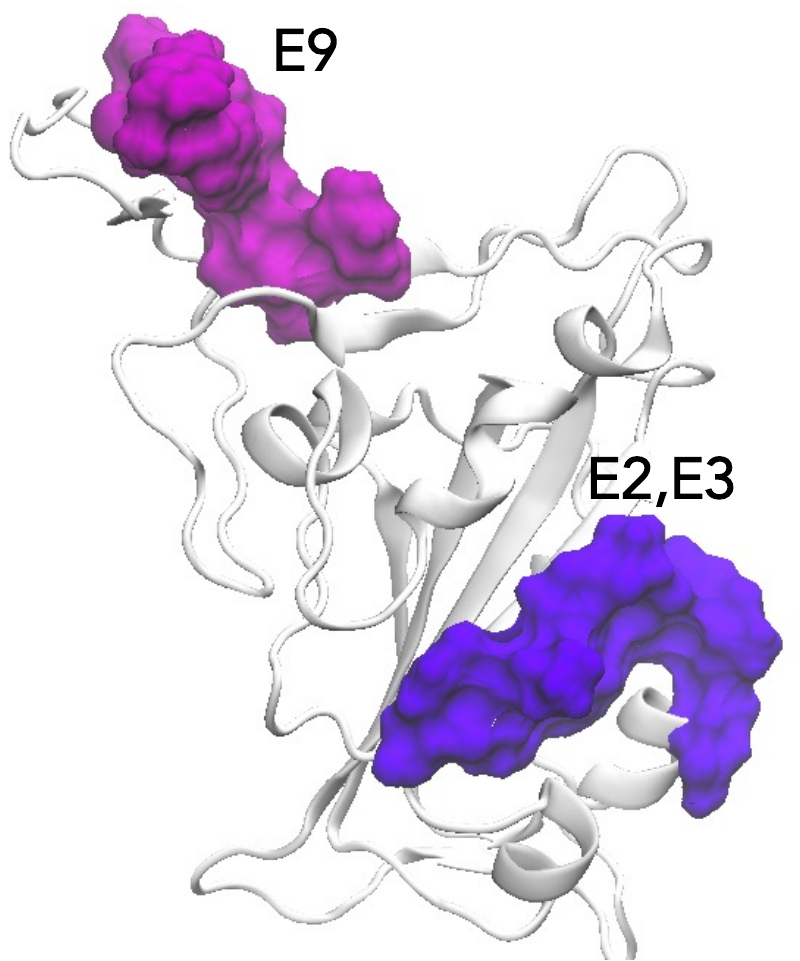

Figure S3. Location of the epitopes E2, E3, and E9 with increased antigenicity in the Omicron RBD. All epitopes have good surface accessibility for the antibody binding.

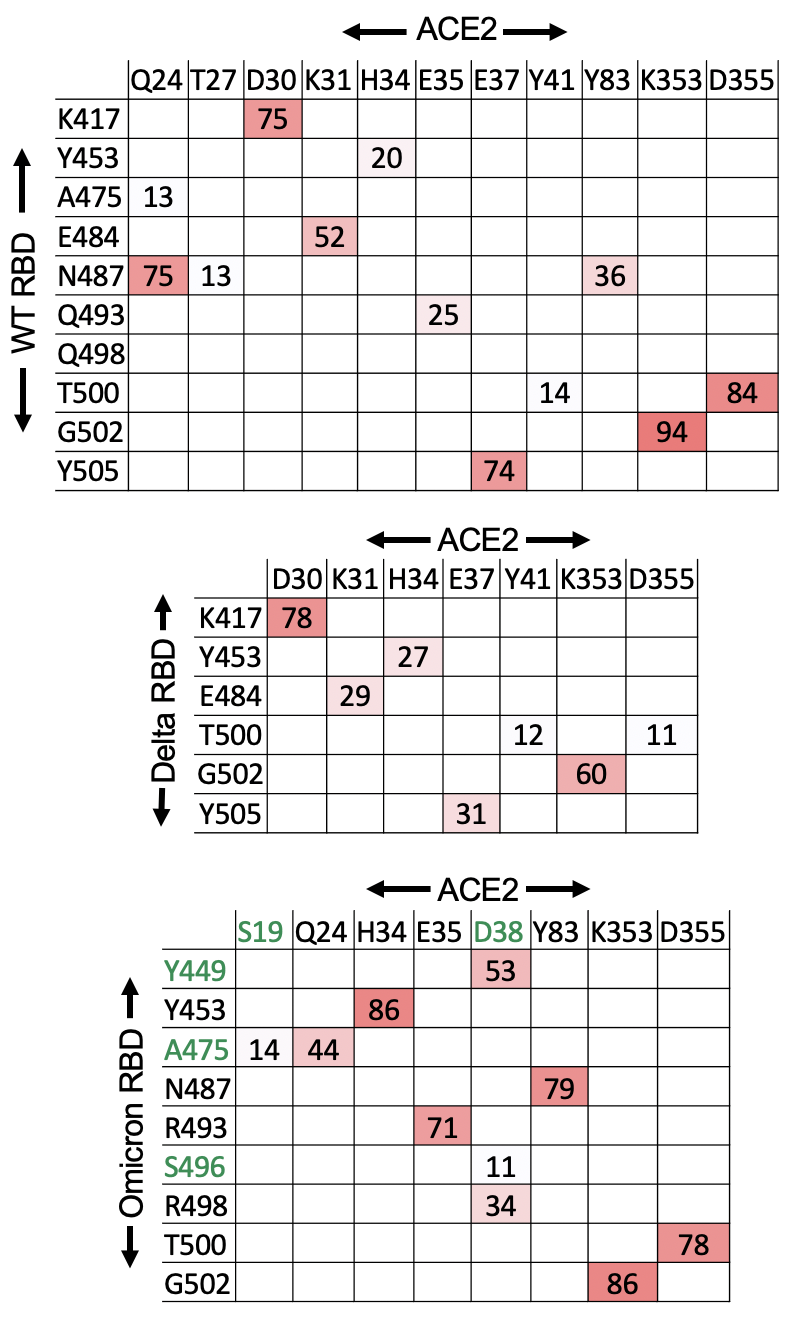

Figure S4. Hydrogen bond pairs for the interprotein interactions between the RBD and ACE2 for the WT, Delta, and Omicron. The matrices for the Delta and Omicron are the same as in Fig. 4a,b and the matrix for Residues participating in the unique hydrogen-bond pairs in Omicron are highlighted in green.

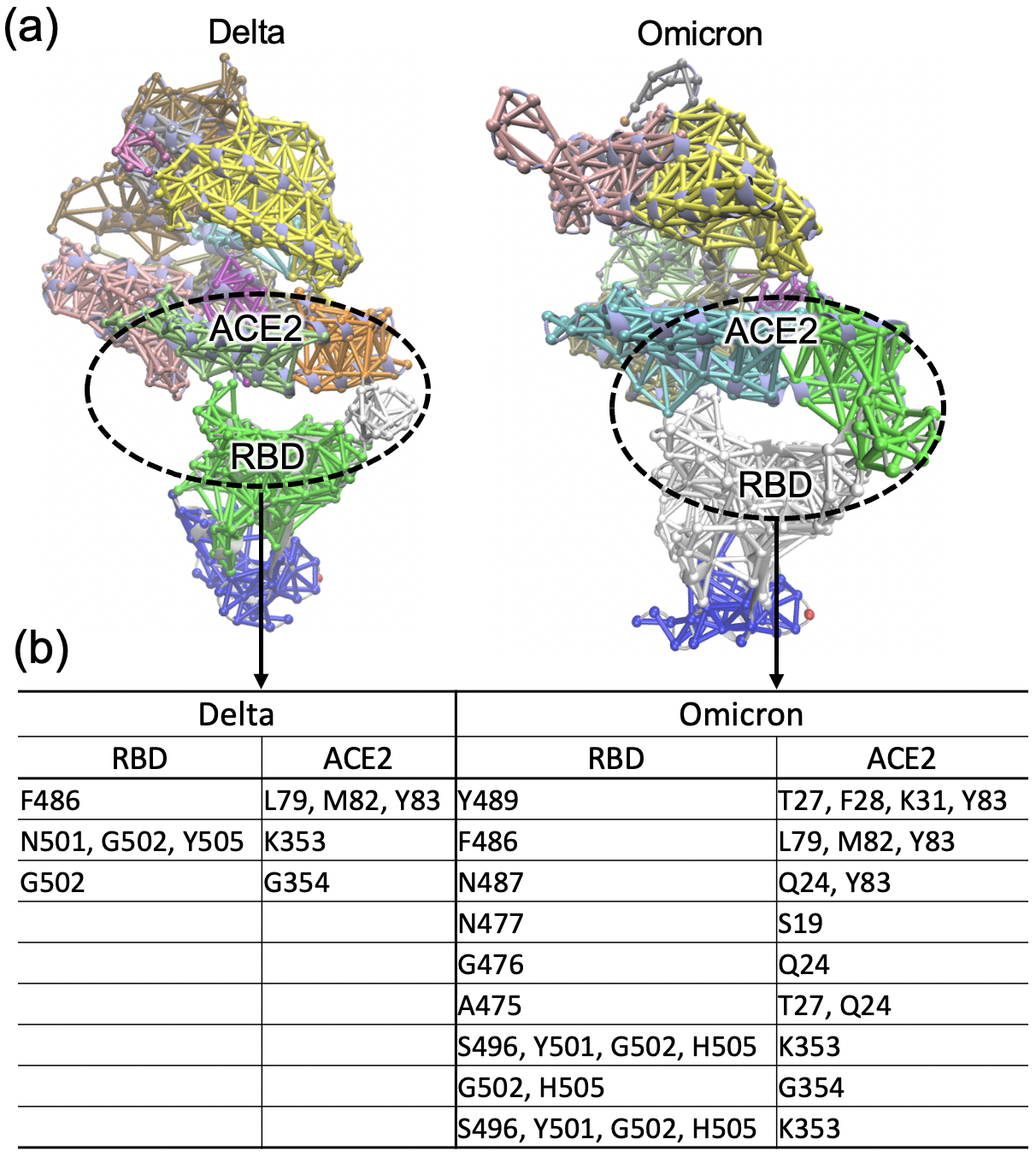

Figure S5. Community analysis performed for the first 50 ns of the trajectories for the Delta and Omicron variants. a) Each identified community is represented by a different color and the region that span both the RBD and ACE2 are circled. b) The amino acid residues involved in the dynamic network of the communities spanning the two proteins (circled region).

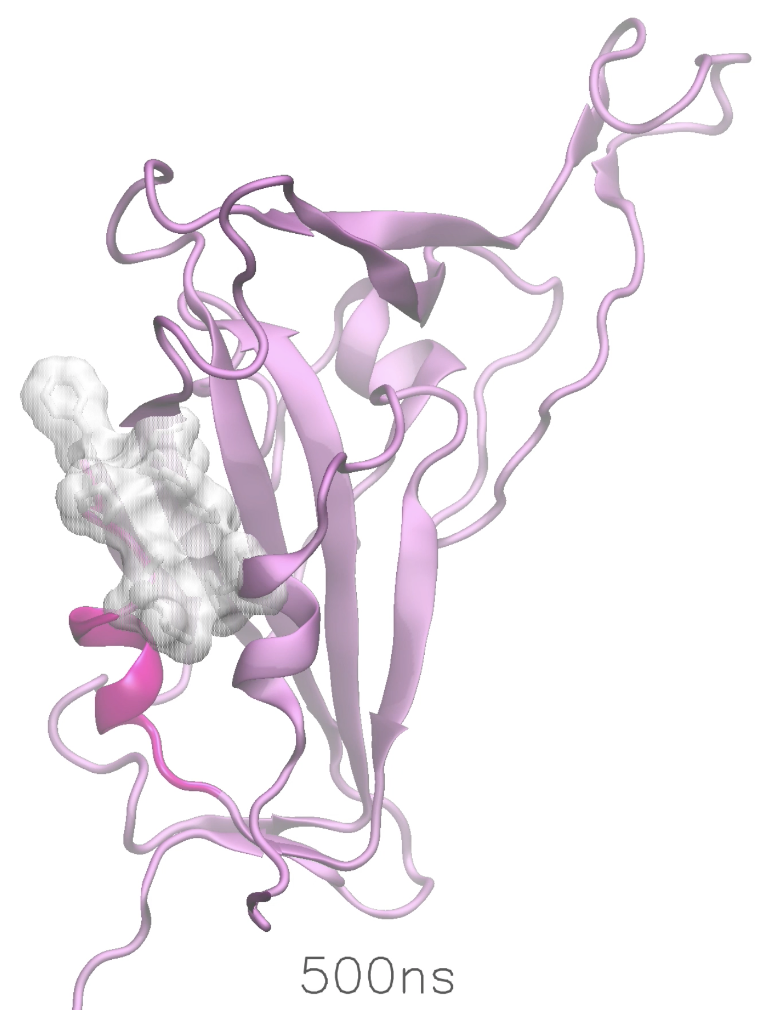

Movie S1. Movie showig the dynamics of the isolated spike RBD of Omicron. The motif comprised of residues 364-375 is shaded in bright purple and the mutated hydrophobic residues L371, P373, and F375 in the motif as well as the nearby hydrophobic residues A435, W436, and F342 are shown in white surface representation.

Table S2. Predicted RBD epitopes that involve mutations in different variants of concern - WT, Alpha (B.1.1.7), Beta (B.1.351), Delta (B.1.617.2), Mu (B.1.621), and Omicron (B.1.1.529) and their corresponding antigenicity values. The mutations in the epitope are underlined. Significantly increased antigenicity in Omicron epitopes E2, E3, and E9 are highlighted in green and a moderately decreased antigenicity in E5 is highlighted in red.

| Epitope | Residues | Variant | Epitope Sequence | Antigenicity |
| --- | --- | --- | --- | --- |
| E1 | 338-356 | WT | FGEVFNATRFASVYAWNRK | 0.24 |
|  |  | B.1.1.529 | FDEVFNATRFASVYAWNRK | 0.28 |
| E2 | 370-378 | WT | NSASFSTFK | 0.12 |
|  |  | B.1.1.529 | NLAPFFTFK | 1.34 |
| E3 | 372-380 | WT | ASFSTFKCY | 0.28 |
|  |  | B.1.1.529 | APFFTFKCY | 1.20 |
| E4 | 417-425 | WT | KIADYNYKL | 1.66 |
|  |  | B.1.351 | NIADYNYKL | 1.55 |
|  |  | B.1.621 | NIADYNYKL | 1.55 |
|  |  | B.1.1.529 | NIADYNYKL | 1.55 |
| E5 | 437-448 | WT | NSNNLDSKVGGN | 0.70 |
|  |  | B.1.1.529 | NSNKLDSKVSGN | 0.24 |
| E6 | 439-451 | WT | NNLDSKVGGNYNY | 0.94 |
|  |  | B.1.1.529 | NKLDSKVSGNYNY | 0.69 |
| E7 | 447-471 | WT | GNYNYLYRLFRKSNLKPFERDISTE | 0.16 |
|  |  | B.1.617.2 | GNYNYRYRLFRKSNLKPFERDISTE | 0.42 |
| E8 | 455-478 | WT | LFRKSNLKPFERDISTEIYQAGST | 0.13 |
|  |  | B.1.617.2 | LFRKSNLKPFERDISTEIYQAGSK | 0.11 |
|  |  | B.1.1.529 | LFRKSNLKPFERDISTEIYQAGNK | 0.10 |
| E9 | 483-493 | WT | VEGFNCYFPLQ | 0.56 |
|  |  | B.1.351 | VKGFNCYFPLQ | 0.60 |
|  |  | B.1.621 | VKGFNCYFPLQ | 0.60 |
|  |  | B.1.1.529 | VAGFNCYFPLR | 1.23 |
| E10 | 491-505 | WT | PLQSYGFQPTNGVGY | 0.34 |
|  |  | B.1.1.7 | PLQSYGFQPTYGVGY | 0.41 |
|  |  | B.1.351 | PLQSYGFQPTYGVGY | 0.41 |
|  |  | B.1.1.529 | PLRSYSFRPTYGVGH | 0.41 |
| E11 | 502-510 | WT | GVGYQPYRV | 1.36 |
|  |  | B.1.1.529 | GVGHQPYRV | 1.01 |

Table S3. MHC-I and MHC-II T-cell epitopes for WT predicted by different methods. The sequences involving mutations in different variants of concern are listed in Table S2.

| MHC-I | | | MHC-II | |
| --- | --- | --- | --- | --- |
| Propred-I | CTLPred | NetCTL | NetMHCII2.3 | EpiTOP3.0 |
| TLDSKTQSL | NLNESLIDL | LTDEMIAQY | FELLHAPAT | VVIKVCEFQ |
| TQSLLIVNN | TRFQTLLAL | WTAGAAAYY | FGAGAALQI | MESEFRVYS |
| FEYVSQPFL | RLDKVEAEV | TSNQVAVLY | VKQLSSNFG | YSKHTPINL |
| GKQGNFKNL | NSPRRARSV | CVADYSVLY | FQTLLALHR | LQPRTFLLK |
| TPINLVRDL | ALDPLSETK | KTSVDCTMY | YYPDKVFRS | YNYKLPDDF |
| VRDLPQGFSALEPL | DLLFNKVTL | STECSNLLL | FLPFFSNVT | VKNKCVNFN |
| QGFSALEPLVDL | AISSVLNDI | GAEHVNNSY | LALHRSYLT | IAARDLICA |
| INITRFQTL | NCDVVIGIV | NIDGYFKIY | LLALHRSYL | MAYRFNGIG |
| FQTLLALHR | KSNIIRGWI | YSSANNCTF | YRLFRKSNL | YRFNGIGVT |
| GTITDAVDCAL | ARDLICAQK | WMESEFRVY | FIAGLIAIV | ILSRLDKVE |
| CYGVSPTKL | TEVPVAIHA | SANNCTFEY | YVGYLQPRT | ITGRLQSLQ |
| IRGDEVRQI | AQKFNGLTV | VASQSIIAY | LIVNNATNV | YVTQQLIRA |
| KIADYNYKL | QSLLIVNNA | NSFTRGVYY | LIANQFNSA |  |
| VVVLSFELL | ATNVVIKVC | CNDPFLGVY |  |  |
| NGLTGTGVLTESNKKFL | YQDVNCTEV | FTNVYADSF |  |  |
| DAVRDPQTL | KNKCVNFNF | GAAAYYVGY |  |  |
| QTQTNSPRR | LFLPFFSNV | RISNCVADY |  |  |
| SIIAYTMSL | FERDISTEI | ERDISTEIY |  |  |
| TNFTISVTT | SVDCTMYIC | ITDAVDCAL |  |  |
| FCTQLNRAL | AYSNNSIAI | RVDFCGKGY |  |  |
| GGFNFSQIL | KRSFIEDLL | NATRFASVY |  |  |
| SKRSFIEDLLFNKVTL | YTNSFTRGV | MTSCCSCLK |  |  |
| FIKQYGDCL | VYAWNRKRI | STQDLFLPF |  |  |
| GDCLGDIAA | VSQPFLMDL | VSNGTHWFV |  |  |
| QKFNGLTVLPPLL | INITRFQTL | GTITSGWTF |  |  |
| IAQYTSALL | IVNNATNVV | RSFIEDLLF |  |  |
| GTITSGWTFGAGAAL | VVFLHVTYV | VLKGVKLHY |  |  |
| AIGKIQDSLSSTASALGKL | TNLCPFGEV | ECSNLLLQY |  |  |
| VVNQNAQALNTLVKQL | ANNCTFEYV | NSASFSTFK |  |  |
| NFGAISSVLNDILSRL | CPFGEVFNA | ASFSTFKCY |  |  |
| QLIRAAEIRASANLA | RVYSTGSNV | QLTPTWRVY |  |  |
| SAPHGVVFL | EILPVSMTK | LSETKCTLK |  |  |
| VYDPLQPELDSFKEEL | SKPSKRSFI | FVFKNIDGY |  |  |
| NHTSPDVDL | GVGYQPYRV | VGGNYNYLY |  |  |
| RLNEVAKNLNESL | FIKQYGDCL | VLPFNDGVY |  |  |
| PWYIWLGFI | GTITSGWTF | FVSNGTHWF |  |  |
| DEDDSEPVL | VIAWNSNNL | YQDVNCTEV |  |  |
|  | QYGDCLGDI |  |  |  |
|  | FSNVTWFHA |  |  |  |
|  | EVFAQVKQI |  |  |  |
|  | NQFNSAIGK |  |  |  |
|  | QTNSPRRAR |  |  |  |
|  | NNATNVVIK |  |  |  |
|  | NATNVVIKV |  |  |  |
|  | QNAQALNTL |  |  |  |
|  | QFNSAIGKI |  |  |  |
|  | WTAGAAAYY |  |  |  |
|  | FVSNGTHWF |  |  |  |
|  | RSVASQSII |  |  |  |
|  | RVVVLSFEL |  |  |  |

Table S4. Sequence based B-cell epitopes predicted by different methods. The sequences involving mutations in different variants of concern are listed in Table S2.

| Bepipred | Bcepred |
| --- | --- |
| QCVNLTTRTQLPPAYTNSFTRGV | MFVFLVLLPLVSSQCVNLTT |
| FSNVTWFHAIHVSGTNGTKRFDN | NLTTRTQLPPAYTNSFTRGVYYPDKVFRSS |
| KSNI | VYYPDKVFRSSVLHSTQDLFLPFFSNV |
| LGVYYHKNNKSWMESEFRVYSSA | SGTNGTKRFDNPV |
| DLEGKQGNFKNLRE | YFASTEKSNIIR |
| HTPINLVRDLPQGFSA | TTLDSKTQSL |
| YLTPGDSSSGWTA | GVYYHKNNKSWMESEFRVY |
| FTVE | MDLEGKQGNFKNLREF |
| YQTSNFRVQP | YFKIYSKHTPIN |
| NITNLC | DPLSETKCTLKSFTVEKGIYQTSNFRVQPTES |
| FGEVFNATRFASVYAWNRK | VYAWNRKRISNC |
| NSASFSTFKCYGVSPTKLNDLCFTNV | RGDEVRQ |
| GDEVRQIAPGQTGKIADYNYK | KIADYNYKLPDDFT |
| NNLDSKVGGNYNY | NSNNLDSKVGGN |
| LFRKSNLKPFERDISTEIYQAGST | GNYNYLYRLFRKSNLKPFERDISTE |
| VEGFNCYFPLQ | CGPKKSTNLVKNK |
| FQPTNG | LTESNKKFLP |
| ELLHAPATVCGPKKSTNLVK | TPGTNTSNQV |
| KKFLP | EHVNNSYECD |
| TNTS | ASYQTQTNSPRRARSVA |
| VNCTEVP | AVEQDKNTQEVF |
| ADQLTPTWRVYSTGSNVFQT | QVKQIYKTPPIKD |
| VNNSYECDIP | LPDPSKPSKRSFIED |
| SYQTQTNSPRRARSVASQS | TLVKQLSS |
| AYTMSLGAENSVAYSN | SRLDKVEAEVQ |
| DKNTQ | LQSLQTYVTQQLI |
| KQIYKTPPIKDFGGF | GQSKRVDFC |
| LPDPSKPSKR | CHDGKAHFPREGV |
| LADAGFIKQYGDCLGD | NNTVYDPLQPELDSFKEELDKYFKNHTSPD |
| VEAEVQI | NIQKEIDRLNEVAKNLNESL |
| GQSKRVDFC | QELGKYEQYIKWP |
| FYEPQIITTD | CKFDEDDSEPV |
| VNNTVYDPLQPELDSFKEELDKYFKNHTSPDVDLGDISGI | EPVLKGVKLHYT |
| ELGKYE |  |
| SCCKFDEDDSEPVLKGVKL |  |

Table S5. B-cell epitopes for WT predicted by Ellipro. The sequences involving mutations in different variants of concern are listed in Table S2.

| Sequence Based | |  |
| --- | --- | --- |
| Residues | Sequence | Score |
| 63-85 | TWFHFDNPVLP | 0.687 |
| 92-105 | FASTNIIRGWI | 0.745 |
| 108-192 | TTLDSKTQSLLIVNNATNVVIKVCEFQFCNDPFLGEFRVYSSANNCTFEYVSQPFLKNLREF | 0.757 |
| 205-218 | SKHTPINLVRDLPQ | 0.678 |
| 236-266 | TRFQTLLALHRGAAAYY | 0.772 |
| 328-381 | RLCPFGEVFNATRFASVYAWNRKRISNCVADYSVLYNSASFSTFKCYG | 0.765 |
| 390-528 | LCFTNVYADSFVIRGDEVRQIAPGQTGKIADYNYKLPDDFTGCVIAWNSNNLDSYNYLYRPLQSYGFQPTVGYQPYRVVVLSFELLHAPATVCGPK | 0.809 |
| 553-564 | TESNKKFLPFQQ | 0.713 |
| 577-585 | RDPQTLEIL | 0.661 |
| 701-717 | AENSVAYSNNSIAIPTN | 0.664 |
| 783-800 | AQVKQIYKTPPIKDFGGF | 0.631 |
| 806-815 | LPDPSKR | 0.552 |
| 883-904 | TSGWTFGAGAALQIPFAMQMAY | 0.623 |
| 1071-1146 | QEKNFTTAPAICHDGKAHFPREGVFVSNGTHWFVTQRNFYEPQIITTDNTFVSGNCDVVIGIVNNTVYDPLQPELD | 0.871 |
| Structure Based | |  |
|  | Residues | Score |
| 1 | A:D1139, A:P1140, A:L1141, A:Q1142, A:P1143, A:E1144, A:L1145, A:D1146 | 0.975 |
| 2 | A:Y707, A:S708, A:N709, A:N710, A:S711, A:I712, A:A713, A:I714, A:P715, A:T716, A:N717, A:Q1071, A:K1073, A:N1074, A:F1075, A:T1076, A:T1077, A:A1078, A:P1079, A:A1080, A:I1081, A:C1082, A:H1083, A:D1084, A:G1085, A:K1086, A:A1087, A:H1088, A:F1089, A:P1090, A:R1091, A:E1092, A:G1093, A:V1094, A:F1095, A:V1096, A:S1097, A:N1098, A:G1099, A:T1100, A:H1101, A:W1102, A:F1103, A:V1104, A:T1105, A:Q1106, A:R1107, A:F1109, A:Y1110, A:E1111, A:P1112, A:Q1113, A:I1114, A:I1115, A:T1116, A:T1117, A:D1118, A:N1119, A:T1120, A:F1121, A:V1122, A:S1123, A:G1124, A:N1125, A:C1126, A:D1127, A:V1128, A:V1129, A:I1130, A:G1131, A:I1132, A:V1133, A:N1134, A:N1135, A:T1136, A:V1137, A:Y1138 | 0.845 |
| 3 | A:L335, A:C336, A:P337, A:F338, A:G339, A:E340, A:V341, A:F342, A:N343, A:A344, A:T345, A:R346, A:F347, A:A348, A:S349, A:V350, A:Y351, A:A352, A:W353, A:N354, A:R355, A:K356, A:R357, A:I358, A:S359, A:N360, A:C361, A:V362, A:A363, A:D364, A:Y365, A:S366, A:V367, A:L368, A:Y369, A:N370, A:S371, A:A372, A:S373, A:F374, A:S375, A:T376, A:F377, A:K378, A:C379, A:Y380, A:L390, A:C391, A:F392, A:T393, A:N394, A:V395, A:Y396, A:A397, A:D398, A:S399, A:F400, A:V401, A:I402, A:R403, A:G404, A:D405, A:E406, A:V407, A:R408, A:Q409, A:I410, A:A411, A:P412, A:G413, A:Q414, A:T415, A:G416, A:K417, A:I418, A:A419, A:D420, A:Y421, A:N422, A:Y423, A:K424, A:L425, A:P426, A:D427, A:D428, A:F429, A:T430, A:G431, A:C432, A:V433, A:I434, A:A435, A:W436, A:N437, A:S438, A:N439, A:N440, A:L441, A:D442, A:S443, A:Y449, A:N450, A:Y451, A:L452, A:Y453, A:R454, A:P491, A:L492, A:Q493, A:S494, A:Y495, A:G496, A:F497, A:Q498, A:P499, A:T500, A:V503, A:G504, A:Y505, A:Q506, A:P507, A:Y508, A:R509, A:V510, A:V511, A:V512, A:L513, A:S514, A:F515, A:E516, A:L517, A:L518, A:H519, A:A520, A:P521, A:A522, A:T523, A:V524, A:C525, A:G526, A:P527, A:K528 | 0.799 |
| 4 | A:F559, A:L560, A:P561, A:F562, A:Q563 | 0.789 |
| 5 | A:F79, A:D80, A:N81, A:P82, A:V83, A:L84, A:P85, A:I100, A:I101, A:R102, A:G103, A:W104, A:I105, A:T108, A:T109, A:L110, A:D111, A:S112, A:K113, A:T114, A:Q115, A:S116, A:L117, A:L118, A:I119, A:V120, A:N121, A:N122, A:A123, A:T124, A:N125, A:V126, A:V127, A:I128, A:K129, A:V130, A:C131, A:E132, A:F133, A:Q134, A:F135, A:C136, A:N137, A:D138, A:P139, A:F140, A:L141, A:G142, A:E156, A:F157, A:R158, A:V159, A:Y160, A:S161, A:S162, A:A163, A:N164, A:N165, A:C166, A:T167, A:F168, A:E169, A:Y170, A:V171, A:S172, A:Q173, A:P174, A:F175, A:L176, A:T236, A:R237, A:F238, A:Q239, A:T240, A:L241, A:L242, A:A243, A:L244, A:H245, A:R246 | 0.756 |
| 6 | A:A27, A:Y28, A:T29, A:R34, A:T63, A:W64, A:F65, A:F92, A:A93, A:S94, A:T95, A:K187, A:N188, A:L189, A:R190, A:E191, A:F192, A:S205, A:K206, A:H207, A:T208, A:P209, A:I210, A:N211, A:L212, A:V213, A:R214, A:D215, A:L216, A:P217, A:Q218, A:S221, A:A222, A:L223, A:G261, A:A262, A:A263, A:A264, A:Y265, A:Y266 | 0.661 |
| 7 | A:G744, A:D745, A:S746, A:T747, A:E748, A:C749, A:S750, A:N751, A:L752, A:L753, A:L754, A:Q755, A:G757, A:S758, A:T761, A:R765 | 0.627 |
| 8 | A:A783, A:Q784, A:V785, A:K786, A:Q787, A:I788, A:Y789, A:K790, A:T791, A:P792, A:P793, A:I794, A:K795, A:D796, A:F797, A:G798, A:G799, A:F800, A:F802, A:Q872, A:A876, A:A879, A:G880, A:T883, A:S884, A:G885, A:W886, A:T887, A:F888, A:G889, A:A890, A:G891, A:A892, A:A893, A:L894, A:Q895, A:I896, A:P897, A:F898, A:A899, A:M900, A:Q901, A:A903, A:Y904, A:N907, A:G910, A:V911, A:T912, A:Q913, A:N914, A:V915, A:L916, A:Y917, A:E918, A:N919, A:Q920, A:L922, A:N925, A:L1034, A:N1108 | 0.61 |
| 9 | A:L984, A:D985, A:P986, A:P987, A:E988, A:E990 | 0.602 |
| 10 | A:R328, A:T531, A:N532, A:L533, A:V534, A:K535, A:N536, A:T553, A:E554, A:S555, A:N556, A:K557, A:Q564, A:R577, A:D578, A:P579, A:Q580, A:T581, A:L582, A:E583, A:I584, A:L585 | 0.597 |
| 11 | A:L806, A:P807, A:D808, A:P809, A:S810, A:K811 | 0.578 |
| 12 | A:A701, A:E702, A:N703, A:S704, A:V705, A:A706 | 0.576 |
| 13 | A:L226, A:V227, A:D228, A:L229 | 0.513 |
| 14 | A:P863, A:L864, A:L865, A:E868, A:M869 | 0.511 |

Table S6. Structure based B-cell epitopes for WT, predicted by Discotope. The RBD residues that are mutated in Omicron are highlighted in boldface and included in Table S2.

| Residue# | Residue | Residue# | Residue | Residue# | Residue |
| --- | --- | --- | --- | --- | --- |
| 281 | GLU | 503 | VAL | 917 | TYR |
| 282 | ASN | 505 | TYR | 918 | GLU |
| 415 | THR | 556 | ASN | 1071 | GLN |
| 420 | ASP | 558 | LYS | 1099 | GLY |
| 449 | TYR | 560 | LEU | 1100 | THR |
| 450 | ASN | 561 | PRO | 1101 | HIS |
| 454 | ARG | 562 | PHE | 1111 | GLU |
| 491 | PRO | 703 | ASN | 1118 | ASP |
| 492 | LEU | 704 | SER | 1140 | PRO |
| 493 | GLN | 705 | VAL | 1141 | LEU |
| 494 | SER | 793 | PRO | 1142 | GLN |
| 496 | GLY | 794 | ILE | 1143 | PRO |
| 498 | GLN | 809 | PRO | 1144 | GLU |
| 499 | PRO | 810 | SER | 1145 | LEU |
| 500 | THR | 914 | ASN | 1146 | ASP |

*References*

20.	Jespersen, M. C.; Peters, B.; Nielsen, M.; Marcatili, P., BepiPred-2.0: improving sequence-based B-cell epitope prediction using conformational epitopes. *Nucleic acids research* **2017,** *45* (W1), W24-W29.

21.	Saha, S.; Raghava, G. P. S. In *BcePred: prediction of continuous B-cell epitopes in antigenic sequences using physico-chemical properties*, International Conference on Artificial Immune Systems, Springer: 2004; pp 197-204.

22.	Kringelum, J. V.; Lundegaard, C.; Lund, O.; Nielsen, M., Reliable B cell epitope predictions: impacts of method development and improved benchmarking. *PLoS computational biology* **2012,** *8* (12), e1002829.

23.	Ponomarenko, J.; Bui, H.-H.; Li, W.; Fusseder, N.; Bourne, P. E.; Sette, A.; Peters, B., ElliPro: a new structure-based tool for the prediction of antibody epitopes. *BMC bioinformatics* **2008,** *9* (1), 1-8.

24.	Irini A Doytchinova, D. R. F., VaxiJen: a server for prediction of protective antigens, tumour antigens and subunit vaccines. *BMC Bioinformatics* **2007**.

25.	Baral, P.; Bhattarai, N.; Hossen, M. L.; Stebliankin, V.; Gerstman, B. S.; Narasimhan, G.; Chapagain, P. P., Mutation-induced changes in the receptor-binding interface of the SARS-CoV-2 Delta variant B. 1.617. 2 and implications for immune evasion. *Biochemical and biophysical research communications* **2021,** *574*, 14-19.

26.	Sztain, T.; Ahn, S.-H.; Bogetti, A. T.; Casalino, L.; Goldsmith, J. A.; Seitz, E.; McCool, R. S.; Kearns, F. L.; Acosta-Reyes, F.; Maji, S., A glycan gate controls opening of the SARS-CoV-2 spike protein. *Nature Chemistry* **2021,** *13* (10), 963-968.
